# Serotonin 5-HT_1A_ receptor binding and self-transcendence in healthy control subjects - a replication study using Bayesian hypothesis testing

**DOI:** 10.1101/226092

**Authors:** Gina Griffioen, Granville James Matheson, Simon Cervenka, Lars Farde, Jacqueline Borg

## Abstract

**Objective:** A putative relationship between markers for the serotonin system and the personality scale self-transcendence (ST) and its subscale spiritual acceptance (SA) has been demonstrated in a previous PET study of 5-HT_1A_ receptor binding in healthy control subjects. The results could however not be replicated in a subsequent PET study at an independent centre. In this study, we performed a replication of our original study in a larger sample using Bayesian hypothesis testing to evaluate relative evidence both for and against this hypothesis.

**Methods:** Regional 5-HT_1A_ receptor binding potential (BP_ND_) was examined in 50 healthy male subjects using PET with the radioligand [^11^C]WAY100635. 5-HT_1A_ availability was calculated using the simplified reference tissue model (SRTM) yielding regional BP_ND_. ST and SA were measured using the Temperament and Character Inventory (TCI) questionnaire. Correlations between ST/SA scores and 5-HT_1A_ BP_ND_ in frontal cortex, hippocampus and raphe nuclei were examined by calculation of default correlation Bayes factors (BFs) and replication BFs.

**Results:** There were no significant correlations between 5-HT_1A_ receptor binding and ST/SA scores. Rather, five of six replication BFs provided moderate to strong evidence for no association between 5-HT_1A_ availability and ST/SA, while the remaining BF provided only weak evidence.

**Conclusion:** We could not replicate our previous findings of an association between 5-HT_1A_ availability and the personality trait ST/SA. Rather, the Bayesian analysis provided evidence for a lack of correlation. Further research should focus on whether other components of the serotonin system may be related to ST or SA. This study also illustrates how Bayesian hypothesis testing allows for greater flexibility and more informative conclusions than traditional p-values, suggesting that this approach may be advantageous for analysis of molecular imaging data.

## Introduction

The serotonin system is involved in a wide range of fundamental physiological functions like regulation of mood, sleep and appetite (Filip and Bader, 2009). Furthermore, serotonergic neurotransmission is implicated in higher brain functions such as cognitive performance (Jenkins et al., 2016) and in several psychiatric disorders, including depression, autism, anxiety disorders and schizophrenia (Fidalgo et al., 2013; Filip and Bader, 2009).

With regard to personality, the serotonin system has been linked to the trait self-transcendence (ST) in both Positron Emission Tomography (PET) and genetic studies (Aoki et al., 2010; Borg et al., 2003; Ham et al., 2004; Kim et al., 2015; Lorenzi et al., 2005; Nilsson et al., 2007; Saiz et al., 2010). ST refers to the degree to which an individual feels part of nature and the universe at large, and to extraordinary experiences such as extra sensory perception and sense of a transcendent being or presence (Gillespie et al., 2003). The association has been interpreted as evidence for a role for the serotonin system in spiritual experiences, as well as providing a putative mechanism for the involvement of serotonin in psychosis, since high scores in ST has been linked to the schizophrenia spectrum disorders (Nitzburg et al., 2014).

Our group previously reported an inverse correlation between 5-HT_1A_ receptor binding potential (BP_ND_), as measured with PET and the radioligand [^11^C]WAY-100635, and ST as measured using Temperament and Character Inventory (TCI). The association was strongest for the subscale spiritual acceptance (SA) (Borg et al., 2003). However, the results could not be replicated in a subsequent PET study at an independent centre (Karlsson et al., 2011). These studies contained 15 and 20 healthy participants, respectively, and therefore, a replication study in a larger sample is required.

### Aims of the study

The aim of the present study was to perform a replication of our original finding of an inverse correlation between 5-HT_1A_ receptor BP_ND_ and ST/SA in a larger sample. In addition to traditional frequentist statistics, we made use of Bayesian hypothesis testing, which allows us not only to test a hypothesis, but also to quantify the relative evidence of the null over the alternative hypothesis. Recently a replication Bayes factor (BF) has been introduced (Verhagen and Wagenmakers, 2014; Wagenmakers et al., 2016), allowing researchers to evaluate replication success by taking the results of previous study into account. In this way, we aimed to evaluate the likelihood of a relationship between 5-HT_1A_ receptor binding and ST/SA from the perspective both of naïve hypothesis testing and of replication success.

## Material and methods

### Subjects

The sample consisted of 50 healthy men: 12 were enrolled as control subjects in a series of different pharmacological studies (for details see Matheson et al. (2015)); 38 in a twin study (Borg et al., 2016). Mean age was 30 ± 5 years (SD). The studies were approved by the Regional Ethics Committee in Stockholm and the Radiation Safety Committee of the Karolinska Hospital, and all subjects provided written informed consent prior to their participation in the studies.

### MR and PET data acquisition (5-HT_1A_ binding potential)

Magnetic Resonance Imaging (MRI) images were acquired using a 1.5TGE Signa system (Milwaukee, WI). T1- and T2-weighted MRI images were acquired for all subjects. The PET system used was Siemens ECAT Exact HR 47 (CTI/Siemens, Knoxville, TN, USA). All subjects were examined using [^11^C]WAY-100635; The injected radioactivity was 276 ± 35 MBq (mean;SD). BP_ND_ values were calculated for the same regions as examined in the original study (Borg et al., 2003): frontal cortex, hippocampus (using the simplified reference tissue model - SRTM) and dorsal raphe nucleus (using a wavelet-based method using the non-invasive Logan plot in order to reduce the noise in this small region). For detailed description see Matheson and co-authors (2015). Other regions were not included in the analysis as they were not part of the original study. However, since [^11^C]WAY100635 BP_ND_ is highly correlated between regions, the inclusion of more regions would therefore be unlikely to provide unique information from the three included regions (Bose et al., 2011).

### Personality assessment

The Swedish translation of the TCI self-report questionnaire was used (Brändström et al., 1998). It consists of 238 true/false items covering four temperament dimensions (novelty seeking, harm avoidance, reward dependence, and persistence) and three character dimensions (self-directedness, cooperativeness, and self-transcendence). Individual scores were calculated for ST and its subscale SA.

### Statistical analysis

Pearson’s correlation coefficients and their corresponding p-values were calculated for the correlation between ST/SA and 5-HT_1A_ BP_ND_ in the frontal cortex, hippocampus and dorsal raphe nucleus. Two BF tests were performed for each comparison. Firstly, we calculated a default correlation BF for the association between BP_ND_ and the ST/SA scores in frontal cortex, hippocampus and dorsal raphe nucleus respectively. Since we specifically wanted to test a negative correlation, we choose a one-sided default Bayes factor test, with a negative Beta prior of width 1 (i.e. flat between −1 and 0) using JASP (JASP Team, 2017). This test compares the predictive adequacy of the null hypothesis H0 (i.e. no correlation) with an alternative hypothesis H- (i.e. a negative correlation). Second, we calculated a replication BF for the correlations for each region as a measure of replication success. This test compares the predictive adequacy of the null hypothesis H0 (i.e. no correlation) with an alternative hypothesis Hr (i.e. original correlation). We slightly modified of the following source code http://www.josineverhagen.com/wp-content/uploads/2013/07/RepfunctionscorrelationFINAL1.txt (for plotting purposes) to the code which can be found online at the following address: https://osf.io/x9gjj/. This code was executed using RStudio (Version 1.0.136) with R 3.3.2 (R Core Team, 2015). We also reanalysed the results of Karlsson et al. (2011) with these methods. BF tests yield a ratio of the relative likelihood of one hypothesis over the other hypothesis, given the data. A BF below 3 indicates weak or anecdotal evidence, > 3 moderate and > 10 strong evidence (Etz and Vandekerckhove, 2016). In this paper, all BFs are presented as the likelihood of the null hypothesis relative to the alternative hypothesis (i.e. BF_0_-specifying a negative correlation as alternative; BF0r specifying the original correlation as alternative). The differences between the default and the replication BF tests can be expressed as follows: the default test addresses the question of whether an effect was present or absent given relatively little prior knowledge of the effect size, while the replication test asks whether the effect was similar to what was found before, or absent (Wagenmakers et al., 2016).

Two potential sources of bias for this analysis were the inclusion of twin pairs, and the use of cerebellar grey matter as the reference region (Hirvonen et al., 2007). We therefore performed two additional analyses by 1) randomly excluding one twin from each twin pair (using www.random.org), resulting in a sample size of 31, and 2) using the white matter as a reference region for hippocampus and frontal cortex.

## Results

In the present sample of 50 subjects, the BP_ND_ of [^11^C]WAY100635 varied about 4-fold between individuals (Table 1). ST scores ranged from 2 to 24 (mean 9.7, SD 5.8); the SA scores ranged from 0 to 12 (mean 3.9, SD 3.1) (Table 1). There were no significant correlations between regional 5-HT_1A_ receptor binding and scores on ST or SA (Figure 1, Table 2).

**Table 1.** TCI scores and BP_ND_ in the original study (Borg et al., 2003) and the present replication study.

|  | Original study |  | Replication |  |
| --- | --- | --- | --- | --- |
|  | Mean (SD) | Range | Mean (SD) | Range |
| TCI scores |  |  |  |  |
| ST | 9.4 (3.8) | 3-15 | 9.7 (5.8) | 2-24 |
| SA | 4.7 (3.0) | 0-9 | 3.9 (3.1) | 0-12 |
| BP<sub>ND</sub> values |  |  |  |  |
| Dorsal raphe nuclei | 2.2 (0.87) | 0.81 - 4.11 | 1.7 (0.48) | 0.64 - 2.88 |
| Hippocampus | 4.7 (1.49) | 1.91 - 7.15 | 5.1 (1.41) | 2.27 - 8.14 |
| Frontal cortex | 3.2 (0.90) | 1.60 - 4.55 | 3.3 (0.73) | 1.21 - 4.61 |

*Abbreviations:* TCI, Temperament and Character Inventory; ST, self-transcendence; SA, spiritual acceptance; BP<sub>ND</sub>, binding potential.

**Figure 1.**
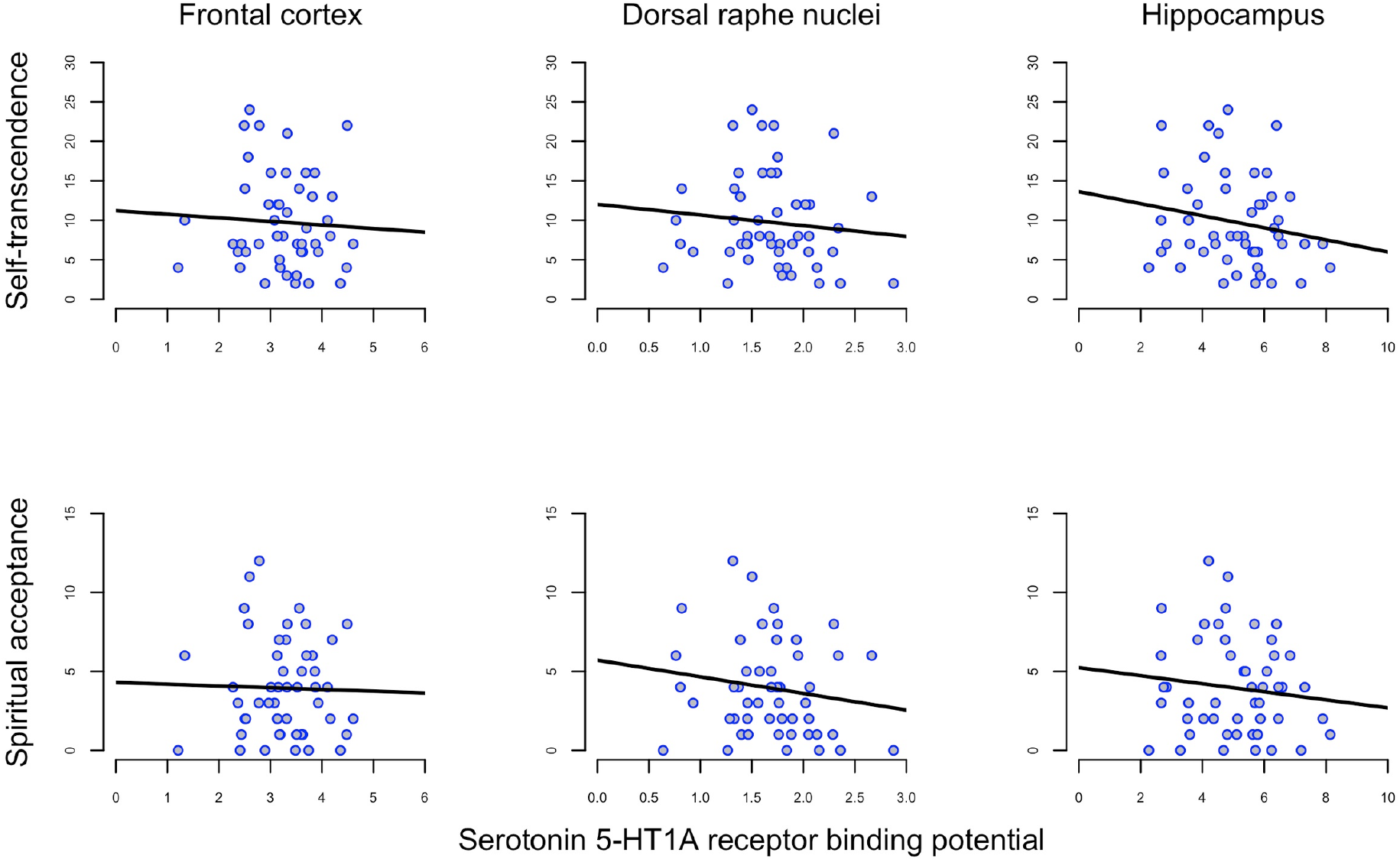
Correlation between self-transcendence (ST) and spiritual acceptance (SA) scales on Temperament and Character Inventory (TCI) and 5-HT_1A_ receptor binding potential (BP_ND_) in frontal cortex, dorsal raphe nuclei and hippocampus in 50 healthy men.

**Table 2.**
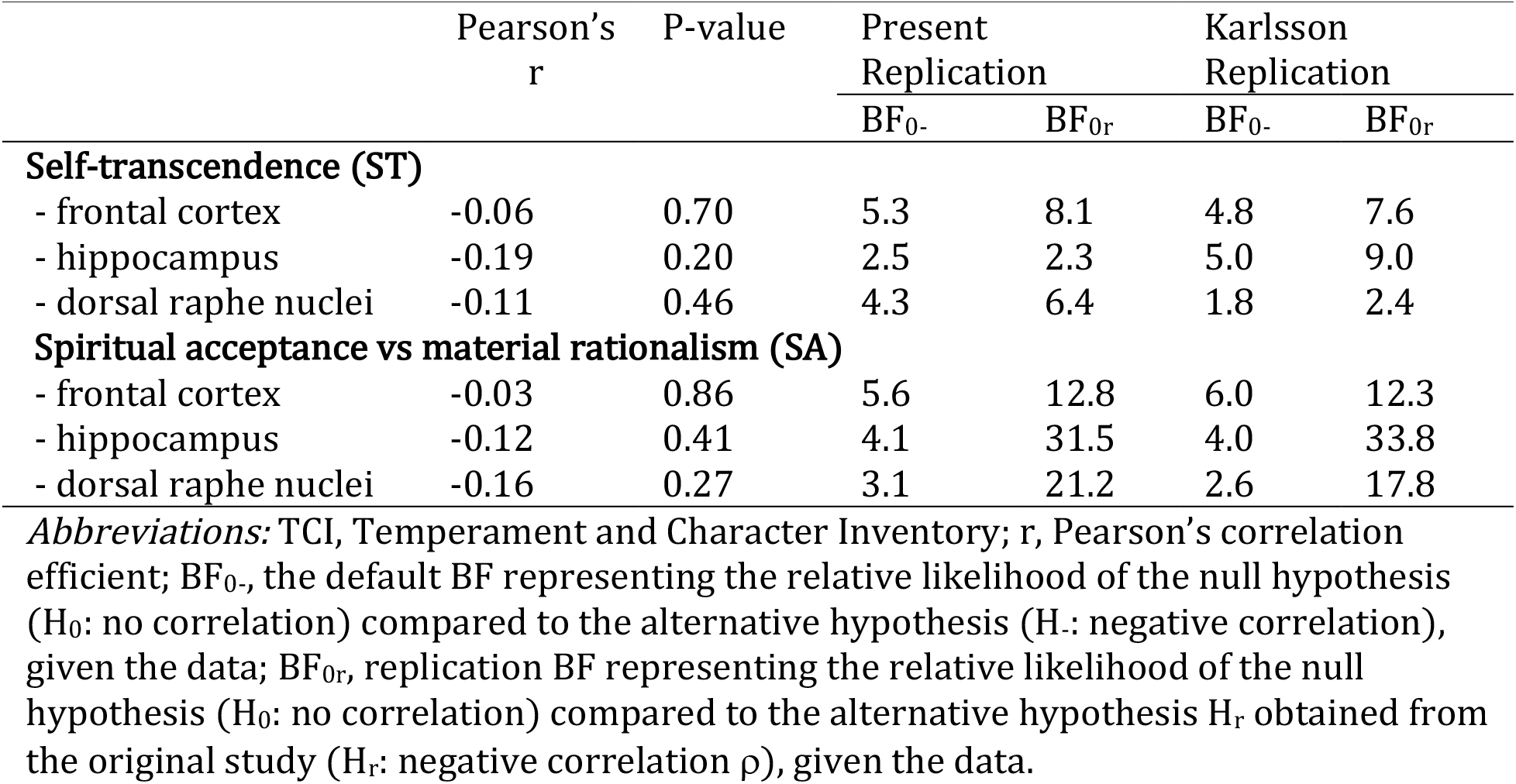
Pearson’s r, default BF and replication BF for 5-HT_1A_ receptor binding and self-transcendence (ST) and spiritual acceptance (SA) assessed by TCI for frontal cortex, hippocampus and dorsal raphe nuclei for present replication and Karlsson’s replication.

**Table 3.**
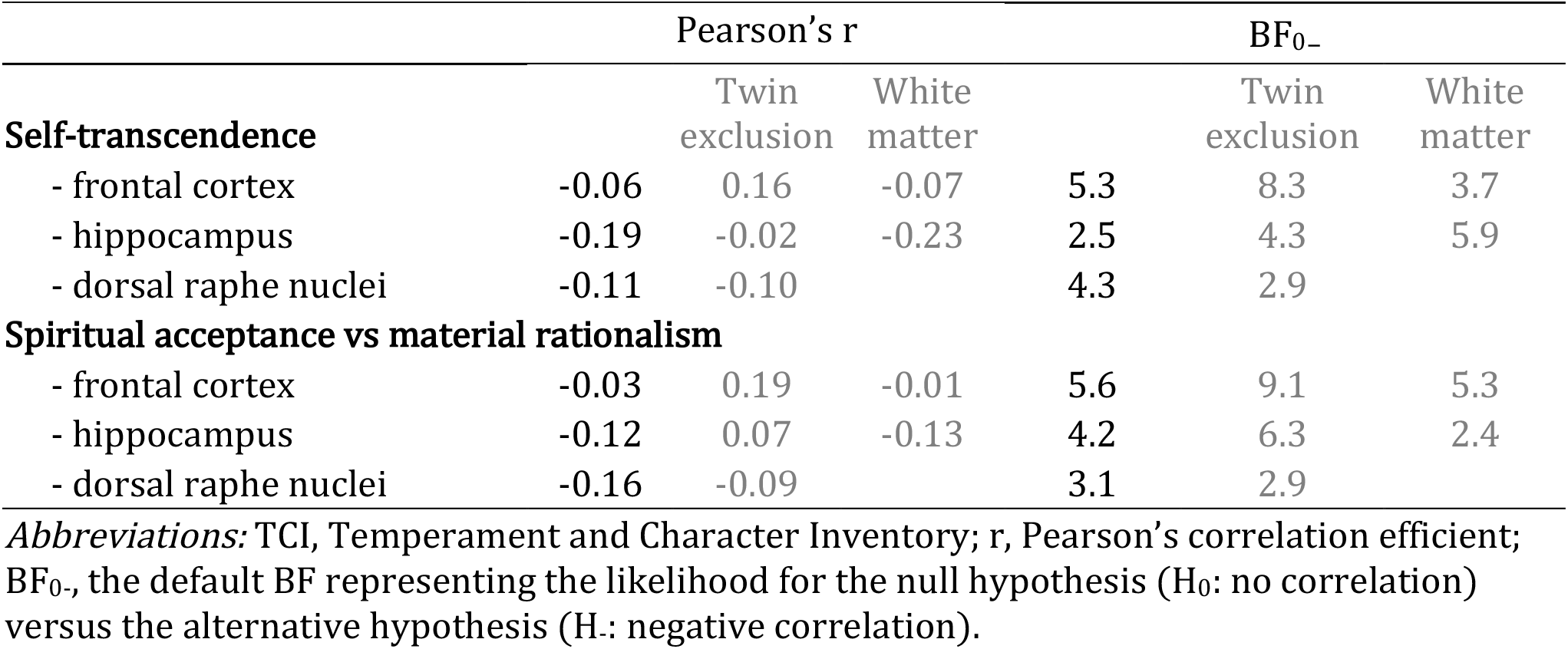
Pearson’s r and default BF for 5-HT_1A_ receptor binding and self-transcendence (ST) and spiritual acceptance (SA) assessed by TCI for frontal cortex, hippocampus and dorsal raphe nuclei for present replication and Karlsson’s replication.

|  | Pearson's r |  |  |  | BF<sub>0-</sub> |  |
| --- | --- | --- | --- | --- | --- | --- |
|  |  | Twin exclusion | White matter |  | Twin exclusion | White matter |
| **Self-transcendence** |  |  |  |  |  |  |
| - frontal cortex | -0.06 | 0.16 | -0.07 | 5.3 | 8.3 | 3.7 |
| - hippocampus | -0.19 | -0.02 | -0.23 | 2.5 | 4.3 | 5.9 |
| - dorsal raphe nuclei | -0.11 | -0.10 |  | 4.3 | 2.9 |  |
| **Spiritual acceptance vs material rationalism** |  |  |  |  |  |  |
| - frontal cortex | -0.03 | 0.19 | -0.01 | 5.6 | 9.1 | 5.3 |
| - hippocampus | -0.12 | 0.07 | -0.13 | 4.2 | 6.3 | 2.4 |
| - dorsal raphe nuclei | -0.16 | -0.09 |  | 3.1 | 2.9 |  |

*Abbreviations:* TCI, Temperament and Character Inventory; r, Pearson's correlation efficient; BF<sub>0-</sub>, the default BF representing the likelihood for the null hypothesis (H<sub>0</sub>: no correlation) versus the alternative hypothesis (H<sub>1</sub>: negative correlation).

Default correlation BFs ranged from 2.5 to 5.6 in favour of the null (Table 2), meaning that the null hypothesis of no correlation is 2.5 to 5.6 times more likely than the alternative hypothesis for a negative correlation. For the results of Karlsson et al. (2011), default correlation BFs ranged from 1.8 to 6.0. 9 out of 12 default BFs provided moderate evidence in favour of the null hypothesis; the remaining 3 provided only weak evidence (Table 2).

The replication BFs ranged from 2.3 to 31.5 in favour of the null hypothesis (Table 2); replication BFs for Karlsson et al. (2011) ranged from 2.4 to 33.8. 10 out of 12 replication BFs provided moderate to strong evidence in favour of the null hypothesis. The remaining 2 replication BFs provided only weak evidence (Table 2).

Figure 2 illustrates the replication BF, showing how the data from the replication study shifts the distribution from the original study towards a correlation coefficient close to zero.

**Figure 2.**
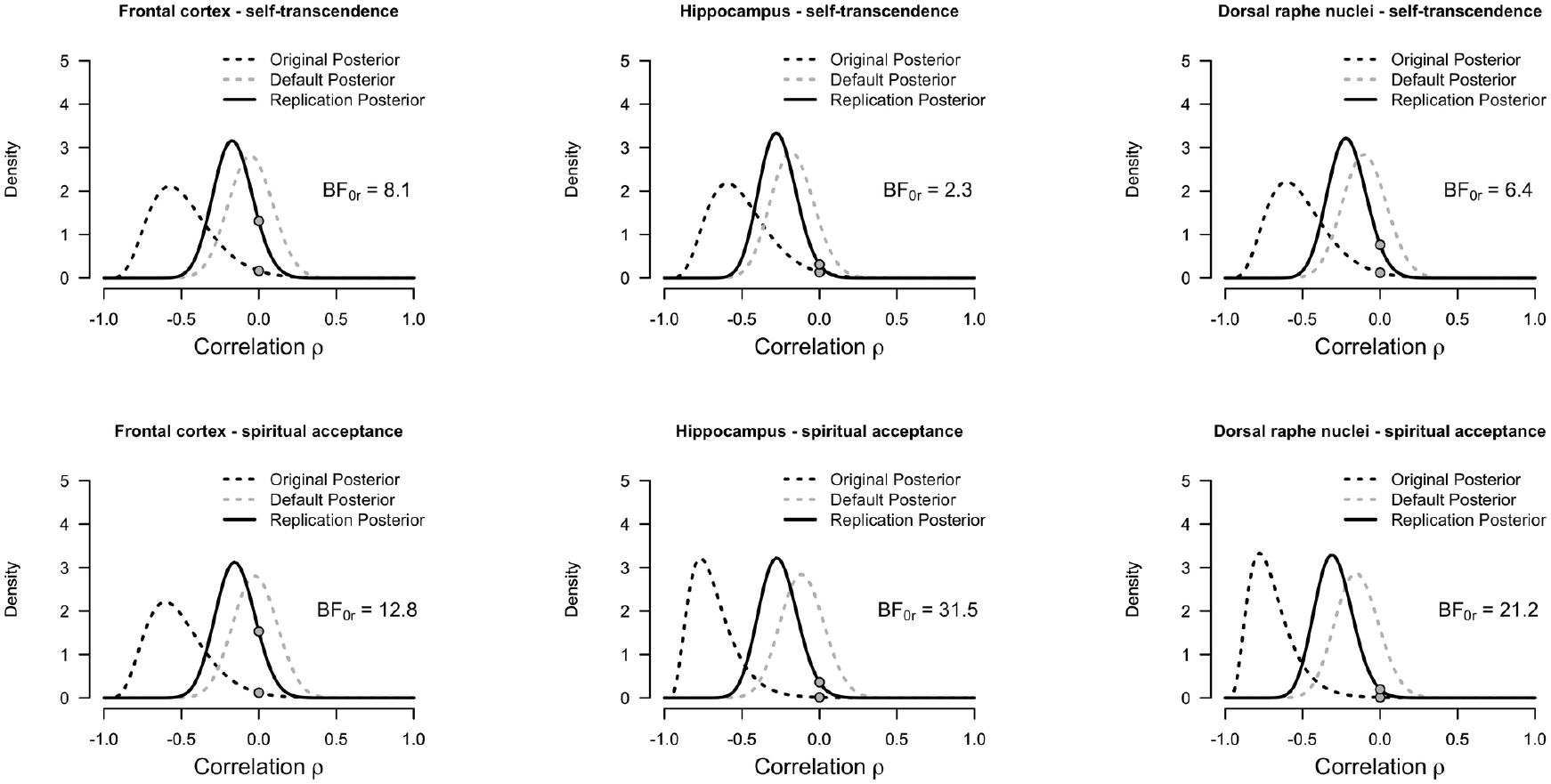
Posterior distributions for correlation between self-transcendence (ST) and spiritual acceptance (SA) on Temperament and Character Inventory (TCI) and 5-HT_1A_ receptor binding potential (BP_ND_) in frontal cortex, dorsal raphe nuclei and hippocampus in 50 healthy men. *The dashed black distribution (named “Original Posterior”) represents the posterior of the original study assuming a flat prior and is used as prior for the replication test. The dashed grey distribution (named “Default Posterior”) represents the posterior of the default test of this replication study (i.e. assuming a flat prior). The black distribution (named “Replication Posterior”) represents the posterior of the replication study using the posterior of the original study (Default Posterior) as a prior. The filled dots represent the height of the prior and posterior distributions for the replication BF calculation at ρ=0. The ratio of these heights yields the Bayes Factor using th e Savage-Dickey density ratio method (Wagenmakers et al., 2010). Abbreviations: BF_0r_, replication BF representing the relative likelihood of the null hypothesis (H_0_: no correlation) compared to the alternative hypothesis H_r_ obtained from the original study (H_r_: negative correlation ρ), given the data.*

The results did not greatly differ after repeating the analysis to account for biases, either by randomly excluding one twin from each twin pair, or by using white matter as reference region (see Supplementary Information).

## Discussion

The aim of the present study was to perform a replication of our previous study (Borg et al., 2003) in a larger sample. We were not able to find any significant relationships between 5-HT_1A_ receptor availability and ST/SA for any of the three regions. This is in line with the results of Karlsson and co-authors in an earlier replication study (Karlsson et al., 2011). Instead, in both this study and in our reanalysis of the results of Karlsson et al. (2011), Bayesian analysis provided more support for the null-hypothesis i.e. that 5-HT1A receptor is not related to the propensity for extraordinary or transcendental experiences

Despite the present results, the serotonin system remains of interest in research on the biological underpinning of personality traits associated with extraordinary experiences. 5-HTT (serotonin transporter) has been linked to ST in both a PET study (Kim et al., 2015), and in genetic studies – though results are conflicting (Aoki et al., 2010; Nilsson et al., 2007; Saiz et al., 2010). Furthermore, 5-HT_1A_, 5-HT_2A_ and 5-HT6 receptor gene polymorphisms have been shown to be correlated to ST (Ham et al., 2004; Lorenzi et al., 2005).

Pharmacological research shows that the serotonin system plays a key role in the effects of hallucinogens, which produce psychosis-like symptoms (comparable to some of the items in the SA scale) (Geyer and Vollenweider, 2008; Vollenweider et al., 1999). Moreover, treatment with SSRI in depressed patients lowered ST scores (Hruby et al., 2009).

Hence, although we failed to replicate the association between 5-HT_1A_ and ST/SA, these lines of evidence motivate further research to clarify the role of serotonin neurotransmission and ST/SA in the healthy population as well as in patients.

The present study was performed on an independent sample of healthy male individuals. Compared to our original study, the sample was more narrow in age range, and 38 of the 50 subjects were twin pairs. TCI scores and BP_ND_ values were however similar to the original study, therefore the more homogenous age range and genetic background of the present sample are unlikely to fully explain the difference in results. Furthermore, we used more advanced image processing methods than in our original study (although many of these, such as automated region of interest (ROI) definition and frame-by-frame realignment of the PET images, were also used in the study by Karlsson and colleagues (Hirvonen et al., 2008; Karlsson et al., 2011)). We were not able to reanalyse the data of the original study using these methods, since T1 weighted MR images were not collected in this sample. However, automated ROIs have been shown to exhibit similar reliability compared to manual (Johansson et al., 2016), suggesting that methodological factors are unlikely to explain the discrepancies.

Replication failure is a common problem in science: in clinical trials and psychology studies replication rates range from 11 to 39%, respectively (Begley and Ellis, 2012; Open Science Collaboration, 2015). Both previous studies on 5-HT_1A_ and ST/SA had low power due to small sample sizes and multiple comparisons without correction, possibly leading to incorrect inferences. According to our calculations using PPV (positive predictive value; the probability that a ‘positive’ research finding reflects a true effect) (Button et al., 2013) the probability that our original finding was true was only around 9%, even before consideration of the two replication studies (see Supplementary Information for the assumptions and the calculation).

### Limitations

Our data consisted of males only. We excluded women from the analysis since the literature is conflicting about the effect of gender and menstrual cycles on 5-HT_1A_ receptor binding (Cidis Meltzer et al., 2001; Costes et al., 2005; Jovanovic et al., 2008; Moses-kolko et al., 2011; Palego et al., 1997; Parsey et al., 2002; Stein et al., 2008; Tauscher et al., 2001) and gender influences ST scores on TCI (Brändström et al., 2001; Garcia-Romeu, 2010). Additionally, we wanted to replicate our original study as closely as possible. Therefore, caution must be exercised when generalizing the present finding in male subjects to the female population. Karlsson and co-authors studied a gender mixed sample (11 males/9 females) in their previous negative study (Karlsson et al., 2011), and in genetic studies, the association between serotonin genes and ST/SA has in some studies been reported to differ between gender (Aoki et al., 2010; Nilsson et al., 2007) whereas others found no difference (Lorenzi et al., 2005; Saiz et al., 2010). As in the original study, we used the cerebellar grey matter as a reference region, which is not considered the gold standard due to small levels of specific binding in this region (Shrestha et al., 2012). However, using arterial plasma to calculate BP_p_ and BP_ND_ using cerebellar white matter as reference, Karlsson and co-authors could not replicate the original findings either (Karlsson et al., 2011). In addition, our analysis using cerebellar white matter showed similar results (see Supplementary Information).

### Strengths

Where Karlsson and co-authors could only conclude that they did not find a significant correlation between ST/SA and 5-HT_1A_ receptor binding (Karlsson et al., 2011), using Bayesian hypothesis testing, we were able to conclude that the data supplied more evidence in favour of the null hypothesis (i.e. no correlation) for both our data and for Karlsson’s results. Furthermore, the replication BF allowed us to take the magnitude of our previous results into account. In this way, using the current data, the replication BF results suggest that the effect reported by the original study was likely either to be overestimated or a false positive; while the default BF results show that the present data is more likely under the null hypothesis given no information about the size of the expected association. As such, these results support the conclusion that there is little to no association between ST/SA and 5-HT_1A_ receptor binding.

Of wider interest in the field of molecular imaging is that Bayesian hypothesis testing provides more informative conclusions than traditional p-values, thus offering pragmatic advantages for analysis of expensive neuroimaging studies, where limited sample sizes are common. For instance, Bayesian hypothesis testing allows for collecting data until the evidence is sufficiently strong to make a conclusion for one or the other hypothesis without requiring correction for sequential analyses. In this way, both costs and radiation exposure can be decreased.

## Conclusions

In conclusion, we failed to replicate our previous finding of a negative association between ST/SA and 5-HT_1A_ receptor binding. Rather, our Bayesian analysis found more evidence for a lack of correlation. Further research should focus on whether other components of the serotonin system may be related to ST/SA.

## Acknowledgements

This study was supported by the Swedish Research Council (2015-02398 (LF); 5232014-3467 (SC)). We gratefully thank the members of the PET group at the Karolinska Institutet, for assistance over the course of the investigation.

## Conflicts of Interest

Conflicts of interest: none. SC has received grant support from AstraZeneca as coinvestigator, and has served as a one off speaker for Roche and Otsuka Pharmaceuticals. LF is employed by AstraZeneca Pharmaceuticals

## Supplementary information

*Results after twin exclusion and using cerebellar white matter as reference region*

### Calculation of PPV

PPV is the positive predictive value (i.e. the chance of the finding being true) and has the following formula:

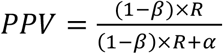

where 1 – β is the power and β the type II error; R is the pre-study odds (in our case the probability of any one personality scale being correlated with 5-HT_1A_ receptor binding); α is the type I error (the statistical significance level).

We made the following assumptions:

- We calculate for an effect size of Pearson’s R = 0.5. This represents a large effect (Cohen, 1988). We calculated the power using this effect size using the “pwr” package in R (Stephane Champely (2015).
- We assume a pre-study odds of 10%. We believe that this is a liberal estimate for comparisons of receptor binding and personality traits.
- The sample size of the original study was 15.
- We set alpha = 0.05 for 10 analyses without correction for multiple comparisons. The original analysis was performed first with three ROIs for seven dimensions, after which self-transcendence was found to be significantly associated with ST (p < 0.05). Due to the high correlations between regional binding estimates, it is an exaggeration to consider this as 21 independent comparisons. Instead, we approximate it as 10 comparisons, comprised of 7 approximately independent original personality dimensions, and three more to account for the multiple ROIs. This means that we assume a final type I error rate of 0.5 (50%).

